# SPANOL (SPectral ANalysis Of Lobes): A spectral clustering framework for individual and group parcellation of cortical surfaces in lobes

**DOI:** 10.1101/203513

**Authors:** Julien Lefèvre, Antonietta Pepe, Jennifer Muscato, Francois De Guio, Nadine Girard, Guillaume Auzias, David Germanaud

## Abstract

Understanding the link between structure, function and development in the brain is a key topic in neuroimaging that benefits from the tremendous progress of multi-modal MRI and its computational analysis. It implies, *inter alia*, to be able to parcellate the brain volume or cortical surface into biologically relevant regions. These parcellations may be inferred from existing atlases (e.g. Desikan) or sets of rules, as would do a neuroanatomist for lobes, but also directly driven from the data (e.g. functional or structural connectivity) with minimum a priori. In the present work, we aimed at using the intrinsic geometric information contained in the eigenfunctions of Laplace-Beltrami Operator to obtain parcellations of the cortical surface based only on its description by triangular meshes. We proposed a framework adapted from spectral clustering, general in scope and suitable for the co-parcellation of a group of subjects. We applied it to a dataset of 62 adults, optimized it and revealed a striking agreement between parcels produced by this unsupervised clustering and Freesurfer lobes (Desikan atlas), which cannot be explained by chance. Already suitable by itself, this spectral analysis of lobes (Spanol) could conveniently be fitted into a multimodal pipeline for optimized and fast lobar segmentation. Eventually, we showed promising results of Spanol on smoother brains and notably on a dataset of 15 fetuses, with an interest for both the understanding of cortical ontogeny and the applicative field of perinatal computational neuroanatomy.

## 1 Introduction

The existence of brain regions associated to specific functions (Friston, 2002) as a dominant paradigm in neuroanatomy is a legacy from the XIXth century. This localizationist view has proved short-sighted and several concurrent segmentations of the brain have been proposed, based on micro architecture first (Zilles and Amunts, 2010), then on neuroimaging studies, in particular with MRI in the past 20 years. *Brain parcellations* have received considerable interest not only because of their deemed underlying biological reality but also because of their advantages in fMRI studies, their ability to reduce the high dimensionality of the data and their use in object-based strategies to overcome the shortcomings of spatial normalization (Thirion et al., 2006). Data-driven MRI based parcellations are most frequently sought, for instance with fMRI data (Thirion et al., 2014) or anatomical connectivity (Lefranc et al., 2016; Parisot et al., 2016) or both (Glasser et al., 2016). Yet brain parcellations based only on the anatomy have been mostly built from information provided by experts and manual labelling. For instance the information of manually delineated cortical sulci has been used to provide volumic (Lohmann and von Cramon, 2000; Tzourio-Mazoyer et al., 2002) or surfacic regions (Cachia et al., 2003; Klein and Tourville, 2012), with different strategies to transfer the labelled information by using registration toward an atlas (Tzourio-Mazoyer et al., 2002) or by working more at the level of the single subject (Cachia et al., 2003; Klein and Tourville, 2012). Even model-based parcellations like the one proposed in (Auzias et al., 2016) require a preliminary labelling of some cortical sulci at the subject level. There are, however, recent attempts of *unsupervised* parcellations procedures, using spectral analysis (Germanaud et al., 2012) or ‘sulcal pits’ concepts (Auzias et al., 2015), the biological correlates of which need further investigations.

Therefore a full unsupervised segmentation of the cortical surface into anatomically meaningful entities remains a very challenging task and there might even be no a priori reasons to hope obtaining one, at least from a non-comprehensive input dataset. Nevertheles in two previous studies of our group (Lefevre et al., 2014; Pepe et al., 2015) preliminary results suggested an intriguing relationships between parcellations obtained by spectral clustering and what is know for a long time as *brain lobes* (Gratiolet, 1854). Brain lobes are a coarse but well-accepted anatomo-functional segmentation of the brain into 5 parts: frontal, parietal, temporal, occipital and insular. This segmentation is not absolute since for instance a 6th limbic part is sometimes proposed. Besides, if unambiguous landmarks ground some lobar limits (e.g. central sulcus as frontal-parietal lateral limit or parieto-occipital sulcus as a parietal-occipital internal limit), others are rather ill-defined (e.g. in the occipital-temporal continuum).

From a general point of view spectral clustering has been largely studied, both theoretically and empirically, mostly in the machine learning community (Ng et al., 2002; Von Luxburg, 2007) but also in computer graphics for mesh segmentation purposes (Liu and Zhang, 2004). Spectral clustering requires a laplacian matrix (in the discrete or graph settings) which corresponds to the Laplace-Beltrami Operator (L.B.O.), often considered as the “swiss army knife” of the geometry processing. Recently this mathematical operator has been more and more popular in the field of neuroimaging, with applications to the description of gyrification pattern (Germanaud et al., 2012; Rabiei et al., 2015), to anatomo-functional variability (Lombaert et al., 2015), in shape registration (Lombaert et al., 2013; Lefèvre and Auzias, 2015) or in shape classification (Lai et al., 2009; Wachinger et al., 2015).

In this article we propose several contributions to both spectral clustering of surfaces with L.B.O and applications to unsupervised, biologically-relevant brain parcellation:

- We describe a *general framework* to parcellate a group of cortical surfaces in an arbitrary number of connected regions based on a spectral clustering algorithm adapted to triangular meshes. Only the geometry of the brain is considered and a limited supervised information (cingulate pole labelling) can be injected to obtain more meaningful results than in (Lefevre et al., 2014).
- Two strategies are investigated to ensure consistent labellings of parcels across individuals: the first one solves the assignment problem for all pairs of segmentations while the second considers a spectral clustering at the groupe level.
- When restricted to 6 regions, the resulting parcellations offer striking *analogies with brain lobes* that can be precisely quantified. The value 6 is optimal regarding a statistical procedure to test the independence of two parcellations.
- The relative robustness of spectral clustering to surface perturbation and in particular unfolding suggests that folding pattern itself play a minor role in the definition of spectral lobes (low frequency description).
- Our spectral clustering approach is also tested on fetal cortical meshes, providing unsupervised segmentations of the immature brain that make sense regarding the adult stage.

## 2 Methods

### 2.1 Spectral clustering

Spectral clustering has become a very popular method in machine learning to segment a set of points in different groups based only on pairwise similarities between those points. A dimension reduction step is performed on a matrix derived from the similarities by extracting its *K* first eigenvectors. Then a classic clustering algorithm such as a *K*-means is performed on these eigenfunctions (Ng et al., 2002; Von Luxburg, 2007).

Graph laplacians are often chosen to define the similarity matrix because of suitable mathematical properties (non-negativity and semi-definiteness) that make them diagonalizable. In our application with cortical surfaces, it is crucial to use a matrix that makes sense in terms of shape and allows fast computations. The Laplace-Beltrami Operator and its discrete representation offer a nice framework because:

- It satisfies the required mathematical properties.
- No parameters tuning is necessary, in contrast to usual similarity graphs that can be obtained with ͛-neighborhood graph, *k*-nearest neighbor graphs or fully connected graph with gaussian weights depending on a *σ* parameter. (Von Luxburg, 2007).
- The derived matrices are sparse which guarantee efficient computation of eigenvectors thanks to Krylov subspace methods as implemented in ARPACK library.
- The resulting eigenvectors correspond to an approximation of the Fourier modes of the ideal cortical surface. They reflect the intrinsic geometry of the brain shape, independently of how it is embedded in the 3D space. From a more physical point of view, Fourier modes correspond to vibration modes of the surface, following a close analogy to the well-known Chladni plates (Chladni, 1787).

Mathematically, Fourier modes correspond to real valued functions *ϕ_i_* defined on a genus-0 surface 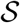 that satisfy:

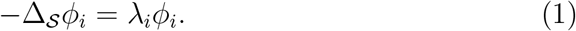

where λ_0_ = 0 ≤ λ_1_ ≤ λ_*i*_… are eigenvalues that can be interpreted as spatial frequencies. 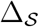 is the Laplace Beltrami Operator of the surface 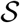 (see (Berger, 2003) for more details).

On a triangular mesh approximating 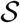 with *N* vertices, it is possible to apply the framework of Finite Element Method. A discretized Fourier mode and its corresponding frequency can be then be obtained as a vector *U* = (*u_i_*)_*i*=1‥*N*_ and a scalar λ such that:

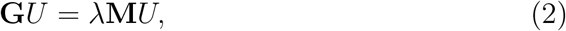

with the stiffness and mass matrices given by :

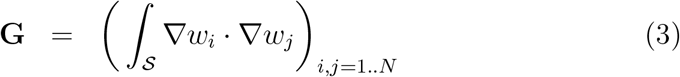

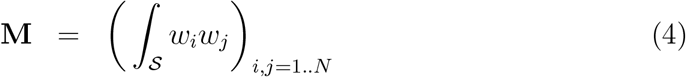

Here *w*_*i*_ is a continuous function, linear on each triangle of the mesh and satisfying the property *w*_*i*_(*i*) = 1 and *w*_*i*_(*j*) = 0 for *j* ≠ *i*. More details on exact values of those matrices can be found for instance in (Lefèvre and Mangin, 2010). The discrete approximation is guaranteed to converge toward the continuous mode when the elementary size of the mesh vanishes (Lai et al., 2009).

The two steps of the spectral clustering adapted to the Laplace-Beltrami Operator are summarized on the left block of Figure 1 and in Algorithm 1.

#### Algorithm 1 Cortical surface segmentation

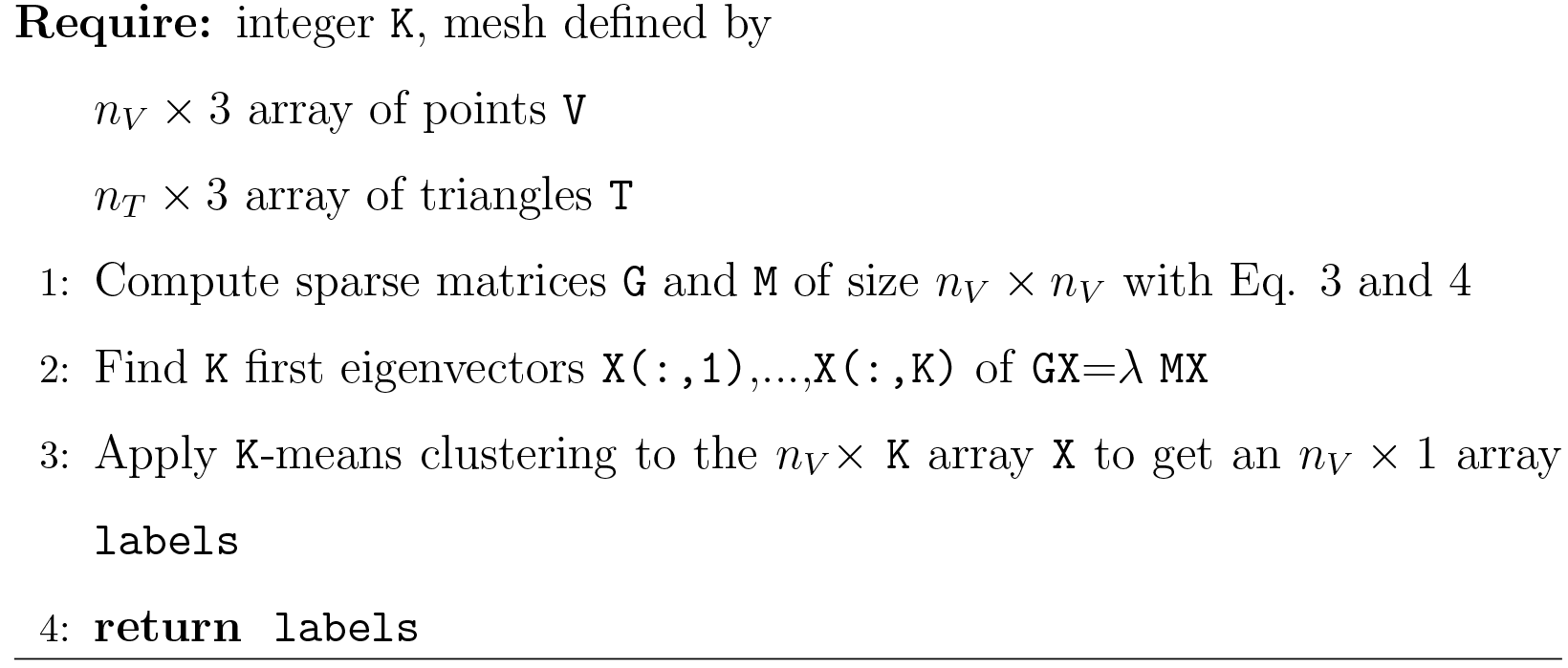

### 2.2 Specific tunings for spectral clustering of lobes

In the previous part we expose the general mechanism to apply spectral clustering on general surfaces. We propose here three variations in the context of cortical surface segmentation.

*Removing of non-cortical vertices in the clustering*. First of all we want to incorporate in the framework some supervised information that would not be efficiently extracted from the data by the spectral analysis. Namely, we consider in our experiments labelled vertices corresponding to the cingular pole, as defined in (Auzias et al., 2013), to be excluded from the spectral clustering. This choice is legitimated by the fact that the surface of the cingular pole cannot be considered as homologous to the rest of the mesh, not being cortical, but rather an artificial filling of the hole created by the segmentation procedure when dissociating the two hemispheres. Futhermore, in line with the underlying hypothesis that Fourier modes may be informative for surface parcellation, it would make sense, if the clustering process had trouble dealing with this artificial surface of cingulate pole, to treat it differently.

From a computational point of view the exclusion of the cingular pole could occur a) before the computation of eigenfunctions, restricting the mesh, or b) before the clustering step, by keeping the eigenfunctions of the entire mesh and excluding labelled points from analysis. Our experiments showed that in the first option, different choices for the boundary conditions (Dirichlet or Neuman) [^2^ Dirichlet boundary conditions for an eigenfunction ϕ correspond to ϕ(*x*) = 0 at each point *x* of the boundary, while Neuman boundary conditions correspond to ∇ϕ(*x*).**n**(*x*) = 0 where **n**(*x*) is a normal vector to the boundary at *x*.] on restricted meshes gave very differents clustering results also very different from processing the entire mesh, suggesting a non-trivial impact on computation. Eventually, we did not find a robust way to deal with this matter. On the contrary, the second option proved to be rather straightforward and labelling of the cingular cluster added interesting regular property to the clustering process. It was thus chosen as a robust way to exclude non cortical surface. In the following we will use the term *constrained spectral clustering* when the cingular pole is excluded of the segmentation process and *unconstrained spectral clustering* in the other case.

*Vary the number of clusters and eigenfunctions*. In the original spectral clustering approach (Ng et al., 2002; Von Luxburg, 2007), the number of clusters equals the number of eigenvectors considered (including the trivial eigenfunction corresponding to eigenvalue 0). This choice is well grounded by linear algebra considerations when one considers a limit case with *K* disconnected regions in a graph whose laplacian matrix has 0 eigenvalue with multiplicity *K* (Von Luxburg, 2007). But the question of using different numbers of eigenvectors and clusters has rarely been addressed and if so in a really empirical approach (Jin et al., 2005).

Here we do not intend to investigate all aspects of this issue but we rather evaluate the qualitative influence of a given Fourier mode on a resulting segmentation. Explicitly, with an anatomical model that includes 6 regions (5 lobes and the cingular pole), the unconstrained clustering requires 6 eigenvectors (including the trivial one) to define 6 clusters and at first sight, the constrained clustering would deal with only 5 eigenvectors to define the 5 unsupervised clusters. Thus, as a test for an add-on effect of a single eigenvector, we will consider only four cases: for the unconstrained approach, the number of eigenvectors is 6 or 7 and 5 or 6 for the constrained clustering.

*Effect of smoothing*. The sensitivity of the spectral clustering to cortical surface deformations may be an important issue both for theoretical (e.g. which aspect of cortical shape impact the clustering) and practical (e.g. stability of the segmentation along development or in case of focal lesion) reasons. We tested this sensitivity by deforming the cortical surface and computing again eigenfunctions and spectral clusters on the deformed surface. For general graphs, this issue of relating clustering error to graph perturbation has been addressed in the litterature of machine learning (Huang et al., 2009). But theoretical results are often limited to two clusters and assumes a small perturbation. We adopted an empirical approach and explored a family of perturbed surfaces obtained by the mean curvature flow (Huisken et al., 1984). This geometric flow is able to smooth a folded cortical surface and it has been shown experimentally that it produces no singularities in the case of cortical surfaces (Lefevre et al., 2013). Moreover, this process is able to mimic the backward emergence of cortical folding pattern (Lefevre et al., 2013). Hence, the combination of mean curvature flow with spectral clustering may be seen as a first step of investigation to ensure the stability of a cortical parcellation along the brain development and its dependency to the folding level

### 2.3. From subject level to group level segmentation

When spectral clustering is performed for different surfaces independently there is no guarantee 1) that clusters exhibit a good reproducibility at the group level and of course, even if segmentation are consistent, 2) that corresponding parcels are assigned to a same label. Starting from our preliminary work (Lefevre et al., 2014) we extended the analysis of point 1) by inspecting extremal parcellations to illustrate the variability. To satisfy point 2) we proposed two different strategies:

- *Matching of individual parcellations* This rellabelling strategy is illustrated on the right block of Figure 1. To achieve consistent labelings, the individual segmentations obtained by the spectral clustering are mapped onto a common spherical template 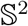 where a relabelling is performed. Note that any spherical mapping approach could be suitable if it transforms the global shape of the brain in a consistent way across subjects. Even ad hoc methods based on affine registration might work in practice but the choice of using a spherical template is suppported by the evaluation of the reproducibility (see later). In this paper we used the Freesurfer method which controls area distortions (Fischl et al., 1999), a property that will be usefull in subsection 2.5. For each surface *s* we have a mapping 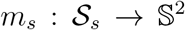. A segmentation map 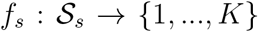 can then be extended on the spherical template as a composition 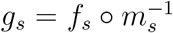. Given two surfaces *s* and *s*′ we can obtain a *K* × *K* table whose value at cell (*k, l*) is given by the cardinality of the set 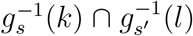. It is simply a quantification of the overlap between two clusters given by labels *k* and *l* for different surfaces. Based on the previous table, it is possible to find a relabelling (i.e. a permutation *σ*_*s*′_ : {1, …,*K*} → {1, …,*K*}) that maximizes the degree of overlap between the two segmentations by taking surface *p* as a reference. This step is achieved via the Munkres algorithm that solves the assignment problem Kuhn (1955).
- *Group spectral clustering* Another possibility consists in applying a global clustering to all the individual eigenvectors pooled together. Formally let us consider surfaces 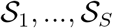 with *N*_1_, …, *N*_*S*_ vertices respectively. **X**_1_, …**X**_*S*_ are the corresponding arrays of *K* first eigenvectors. We can build a big vector **X** in ℝ^*N × K*^, where *N* = *N*_1_ +… + *N*_*S*_, as a concatenation of all the eigenvectors of the *S* surfaces. The *K*-means clustering is applied to the vector **X** which yields a big vector of labels that produces individual segmentations. This step is summarized in Algorithm 2 and on Fig 2. As noted on Fig 2, a technical step is required for each eigenfunction at the beginning of the process. Given an eigenfunction Φ associated to an eigenvalue λ, note that —Φ is also an eigenfunction. To overcome this sign ambiguity issue, we used a simple correlation of two eigenfunctions resampled on the common sphere and switched the sign of one eigenfunction if the correlation was negative.

**Figure 1:**
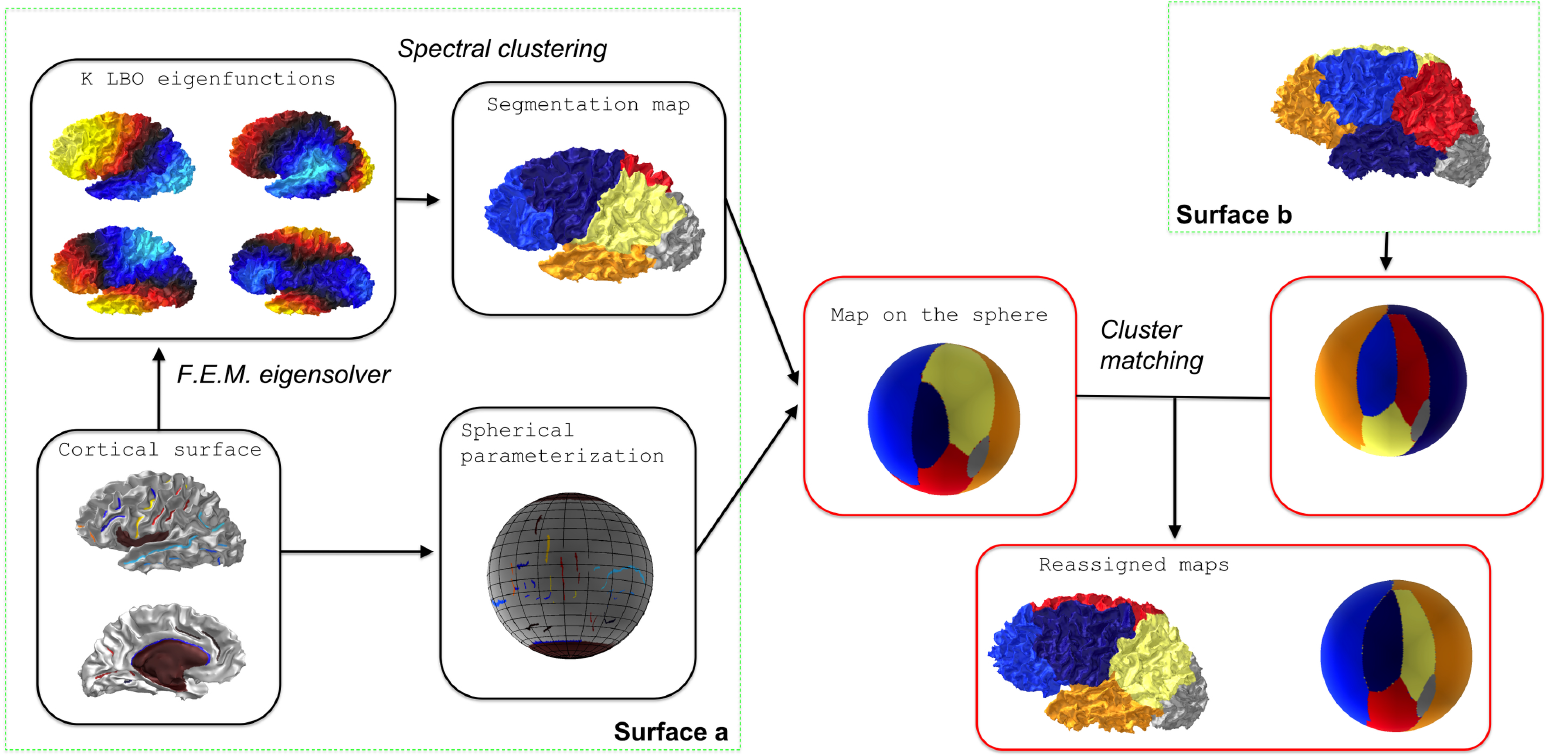
Flowchart of individual spectral clustering and inter-individual strategy for relabelling. The cluster matching step is achieved via the Munkres algorithm.

**Figure 2:**
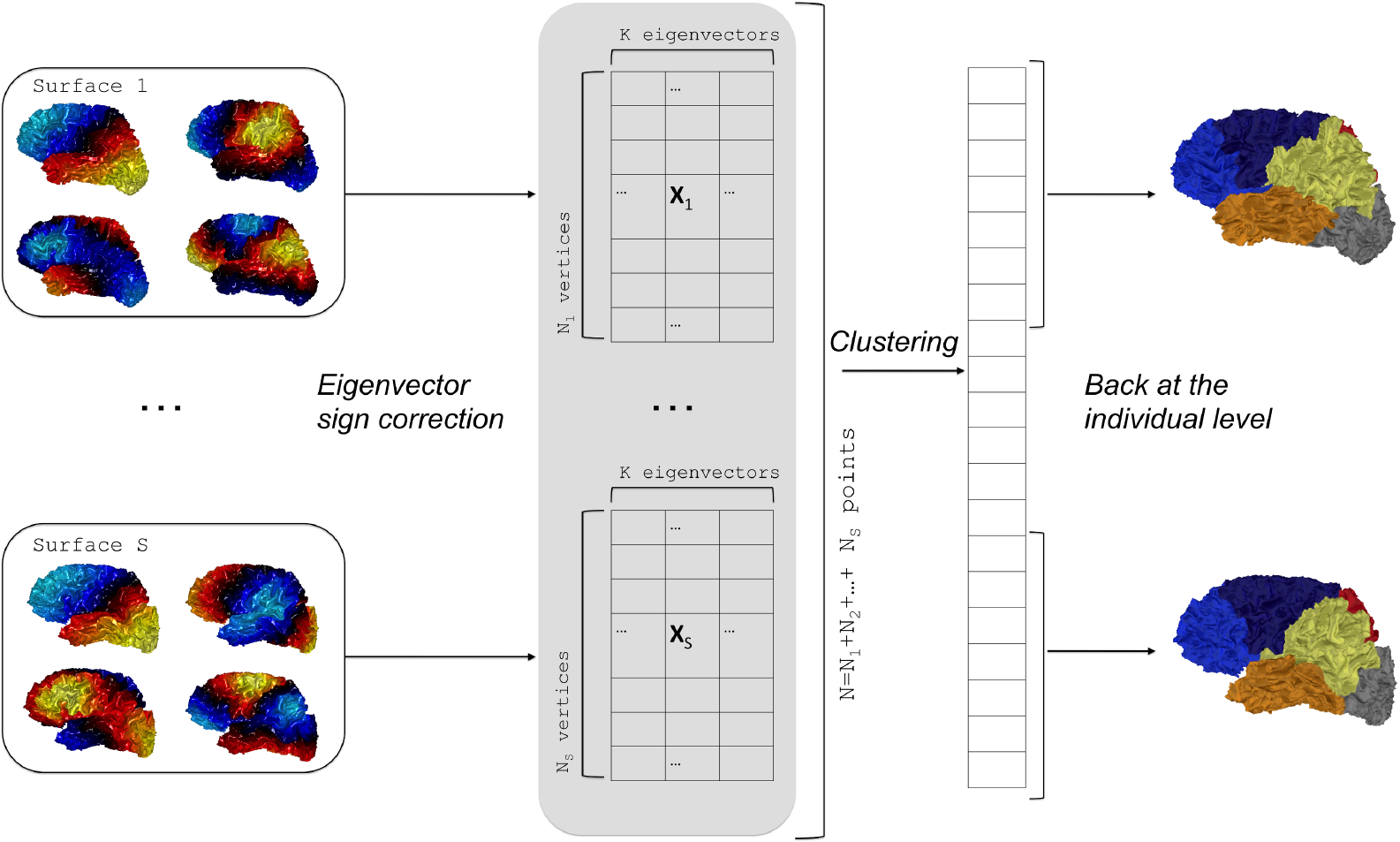
Flowchart of group spectral clustering. Individual eigenvectors are computed for each surface and the sign ambiguity is corrected. Concatenation of eigenvectors yields a large array on which a clustering is performed. Last the labels are back-projected on the individual surfaces.

#### Algorithm 2 Group spectral clustering

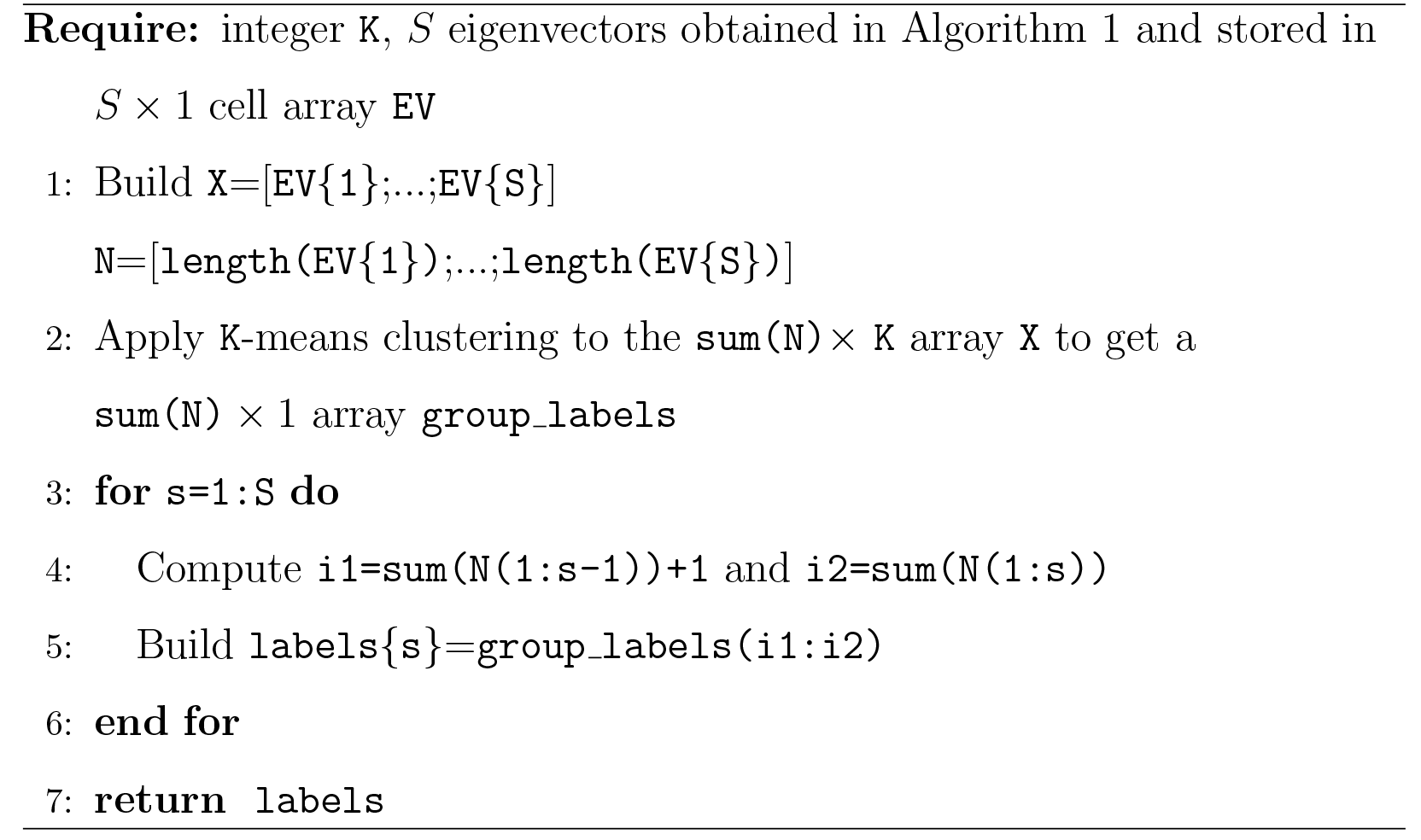

### 2.4 Evaluation metrics

We propose here different metrics to evaluate the quality of parcellations across a group of surfaces. The first one assesses the relationship between cluster boundaries and some anatomical landmarks that have been identified by neuroanatomical experts (sulcal lines). The second metric quantifies the distance between spectral clusters and reference clusters (obtained by concatenating smaller parcels obtained with the Desikan atlas in the Freesurfer software (Desikan et al., 2006)).

*Spectral boundaries and landmarks*. We consider *M* anatomical landmarks that will be compared to cluster boundaries (*M* = 2 in practice). In particular, Central Sulcus (C.S.) and Parieto-Occipital Sulcus (P.O.S.) are very important landmarks since they are reliable borders between lobes. For each discrete sulcal line 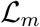 we can define a mean distance to the boundaries of the segmentation map 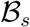 of subject *s*:

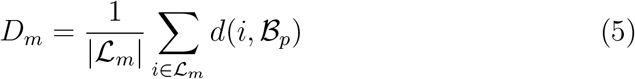

where *d*(.,.) is the geodesic distance from a point to a set of points. In practice we use the Fast Marching Algorithm and the Matlab implementation of (Peyré and Cohen, 2008) to obtain a distance map from 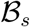 to any vertex of the mesh.

*Comparison of two parcellations*. One can compare two segmentations of a mesh by considering them as partitions of the *n* vertices of the template sphere and computing a distance between these partitions. Note that the two parcellation maps may have different number of labels and we will denote them as *K* and *L* in the following.

We have used the *rand distance* which is frequently employed to obtain partition distances or partition similarities (*rand index*). It has the interesting property to be a mathematical distance. The very fast computation time is a clear benefit over distances such as the transfer one (Denœud and Guénoche, 2006) when it comes to compute extensively pair-wise distances on groups of segmentations.

The rand distance is based on the number *a* of pairs of vertices that are in the same cluster in the two partitions, and the number *b* of pairs of vertices that are in different clusters in both partitions. The values of *a* and *b* can be obtained from the contingency table between the two segmentations *T_k,l_* (e.g. following (Denœud and Guénoche, 2006)):

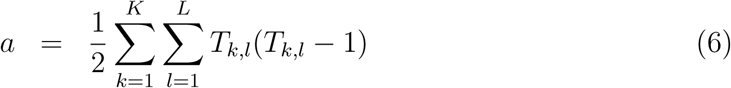

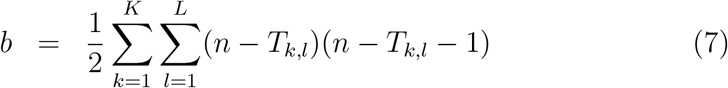

Then the formula for the rand distance is simply :

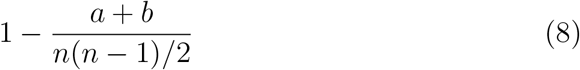

To evaluate the overlap between each Freesurfer lobe and the corresponding spectral cluster we have chosen *Dice coefficients*, which are commonly used in neuroimaging but do not correspond to mathematical distances. The Dice between two sets *A* and *B* is given by:

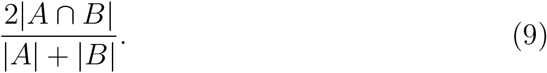

0 corresponds to no overlap while 1 implies equality of *A* and *B*.

*Consensus parcellation*. The problem of obtaining a meta-clustering that summarizes several individual clusterings comprises a vast litterature with different terminology, from “consensus clustering” (Strehl and Ghosh, 2002) to “consensus of partitions” (Guénoche, 2011).

In our case we aim at obtaining consensus maps for visualization purposes and we have used fast and simple methods.

First it is possible to define an intuitive average segmentation in *K* clusters by a *voting strategy*. At each vertex *i* of the sphere we consider the label that is present the most which can also be mathematically written as:

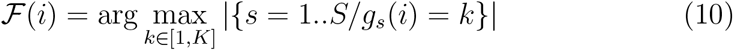

Another possibility is to compute the *median parcellation* based on the *S* × *S* distances between all the segmentation maps of the *S* subjects.

In practice the two resulting segmentations were very similar and we adopted simply the consensus map obtained by the voting strategy in the following.

### 2.5. Statistical evaluation

We propose a statistical test to determine if a brain parcellation (a mapping *f_s_*) obtained by spectral clustering has a statistical relationship with a segmentation map obtained by an expert system, e.g. a Freesurfer parcellation.

For a given subject *s*, an expert system provides a mapping 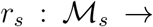 {1,…, *L*}. Then a positive value can be obtained by evaluating *Dist*(*r*_*s*_, *f*_*s*_), a distance between the ground truth *r_s_* and the reference segmentation *f_s_*. This value by itself does not reflect something obvious to interpret and we need to compare it to a distribution of values under a null hypothesis of independence between the reference and the automatic segmentation. This distribution can be generated by randomly rotating the maps *g_s_* defined on the sphere, provided that the sherical parameterization *m_s_* preserves areas as much as possible to avoid some biases in the spatial distribution of random parcellations on the original mesh 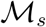. Eventually the distribution yields a p-value.

To obtain random rotations, we generate 3 × 3 matrix with coefficients following independant normal distributions, apply a *QR* decomposition and keep the orthogonal matrices *Q* which are uniformly distributed (Blaser and Fryzlewicz, 2016).

We can sum up the procedure in the following algorithm and in Fig 3.

**Figure 3:**
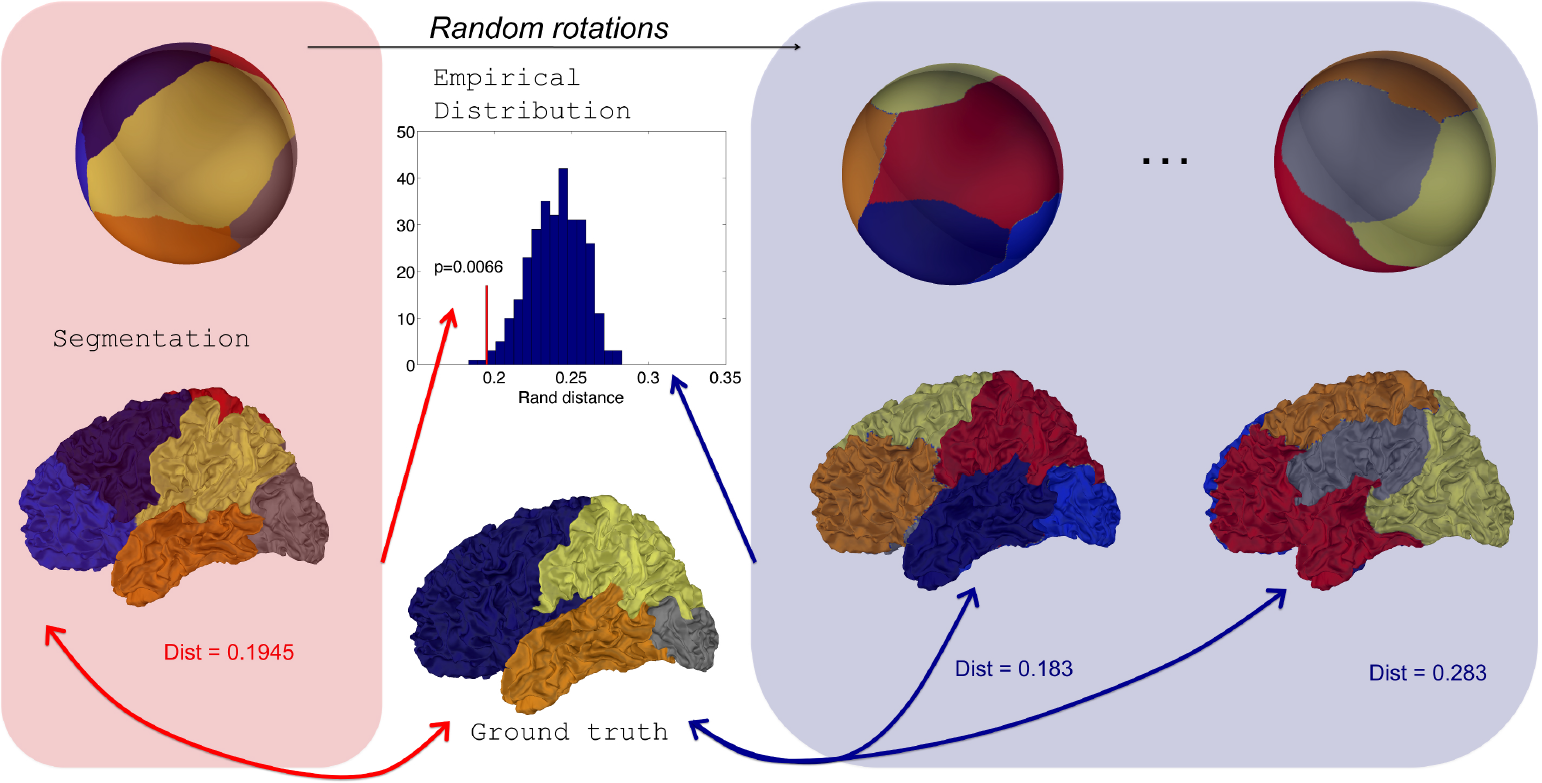
Distribution of Rand distances obtained through random rotations of a parcellation on the sphere. From an initial segmentation projected onto a regularly sampled sphere, random rotations are applied to generate random parcellations onto the sphere and consequently on the original surface. Rand distances are evaluated for each random parcellation which yields an empirical distribution.

#### Algorithm 3 Signicance of Rand Distance

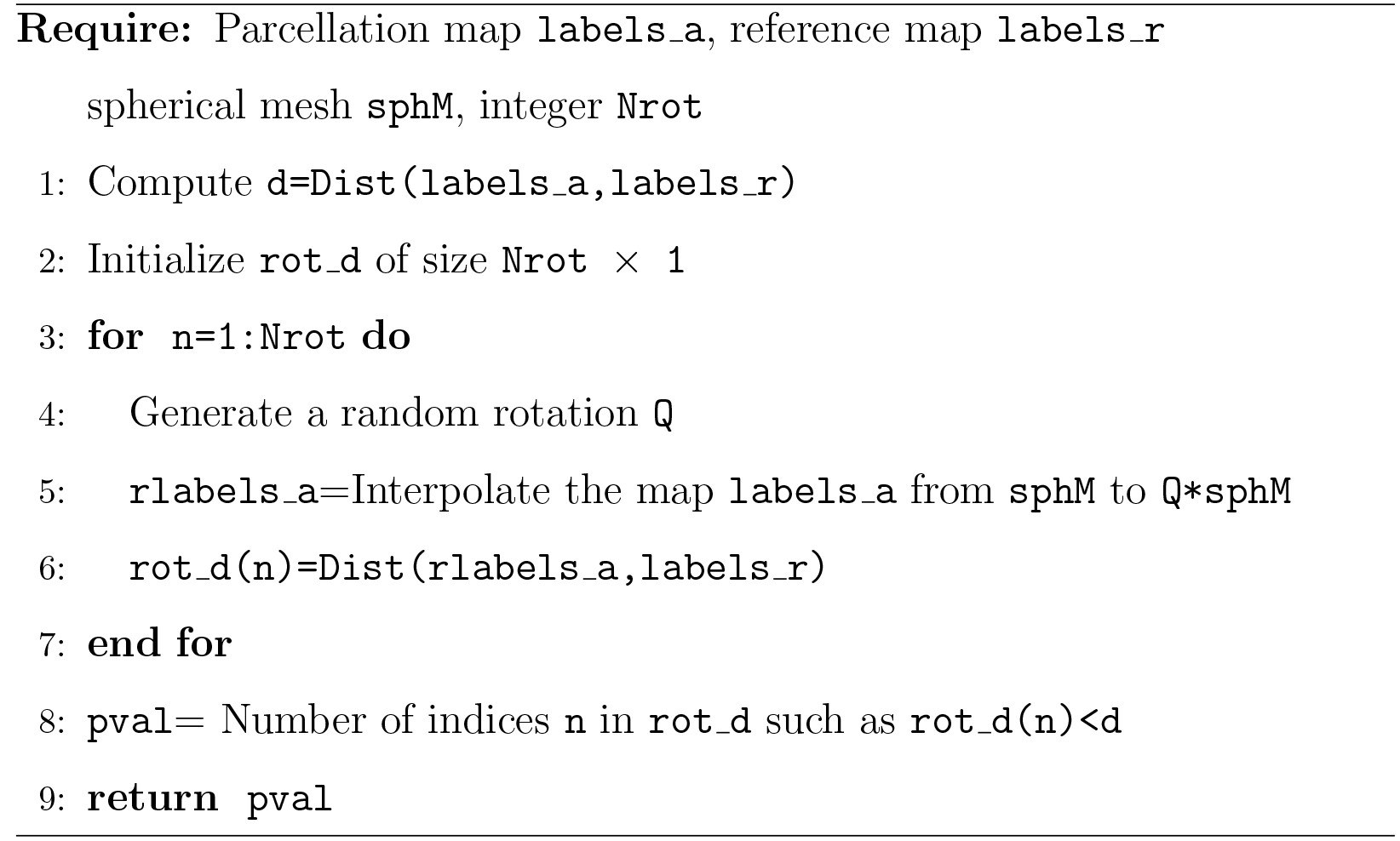

## 3. Results

### 3.1. Datasets and pre-processings

First we tested our method on a dataset of 62 left hemispheres previously used in (Auzias et al., 2013). Triangular meshes were obtained from T1-MRI through the morphologist pipeline of BrainVisa software. Central and Parieto-Occipital sulci were delineated semi-automatically by using Surf-paint (Le Troter et al., 2011). In parallel, Freesurfer was used to obtain 1) a spherical parameterization of each mesh and 2) to generate a parcellation of each surface in 35 regions based on the atlas of (Desikan et al., 2006). The 35 regions were then merged in 4 lobes (frontal, parietal, temporal, occipital), insula and cingulate pole. Because our point in this study was not to discuss in the details the position of the the lobar limits that are not as grounded as the central sulcus of the parieto-occipital one, we kept with the merging proposed on Freesurfer wiki [^3^ Cortical Parcellation/Lobe Mapping: https://surfer.nmr.mgh.harvard.edu/fswiki/CorticalParcellation] without further refinements.

We also tested the constrained spectral clustering on a dataset of 15 fetuses previously used in (Lefèvre et al., 2015). The gestational age ranges from 21 weeks to 34weeks. Triangular meshes were obtained from T2-MRI and preprocessed with methods dedicated to fetal imaging. Major sulci, when present, were traced with Surfpaint.

Next the results are divided in four subsections. In subsections 3.2, 3.3, 3.4 results involve the first dataset. Eventually the last subsection illustrated some stability properties of the spectral partitioning up to surface deformation and how it can be a powerful tool to parcellate developing brains as revealed by the second dataset.

### 3.2. Descriptive analysis of spectral clusters

On Fig 4, we show consensus parcellations obtained with the unconstrained (top) and constrained (bottom) approaches (*K* = 6 in both cases) in parallel with a consensus parcellation in lobes obtained with Freesurfer (middle).

The comparison of unconstrained clustering and FreeSurfer reference (first two rows) showed that spectral clusters were strongly reminiscent of anatomical lobes, even with a plain unrefined strategy. Indeed there were several differences between segmentations: a) the insular lobe was not segmented and was divided by frontal, parietal and temporal lobes, b) the frontal lobe was divided in two parts (light and dark blue) suggestive of prefrontal and precentral subdivisions, c) the cingulate pole was poorly delimited and even included in a large mesial region (in red) which extended on a short portion of the external face. This parcel had little anatomical correlate and clearly impaired mesial segmentation. In this view, the introduction of a constrained cingulate pole resulted in several qualitative improvements on the consensus map (third row of Figure 4) that were also present at the individual level on Figure 6. By definition of the method, the cingulate pole was perfectly segmented, but it resulted in a global improvement of mesial segmentation, particularly the mesial part of the parietal lobe that was then much more consistent with the one by Freesurfer (second row) while the parietal-occipital boundary remained almost unchanged. Interestingly, the better accordance with Freesurfer segmentation extended on the external face where the temporal-occipital boundary was shifted more posteriorly. Other lobes did not show any important changes.

In details, the most striking similarities between spectral segmentation and reference correspond to the two boundaries, between frontal and parietal lobes (dark blue, yellow) and parietal and occipital lobe (gray, yellow). At the individual level we obtained three reference subjects for which the distances between central sulcus (resp parieto-occipital sulcus) and the corresponding boundary were the smallest, the median and the largest (see Fig 5). Another look on the variability with three extreme cases was also provided on Figure 6 and Figure 1SI in Supplementary Information by using another metric based on comparisons with Freesurfer lobes (see subsection 3.3 for quantitative aspects).

### 3.3. Quantitative validation of optimal spectral segmentation

After this visual description, we proposed some quantitative results to emphasize striking similarities between spectral regions and brain lobes. We will now consider mostly the constrained strategy with *K* = 6. In Supplementary Information we added other quantitative results regarding the different clustering strategies (constrained/unconstrained).

*Region boundaries and sulci*. First we obtained distances between two well-defined boundaries of lobes and corresponding spectral boundaries. Based on our definition 5, we computed two distances *D*_1_ and *D*_2_ for the Central Sulcus (C.S.) and the Parieto-Occipital Sulcus (P.O.S.) of each subject. The numeric values provided a first view on the variability of the segmentations. On Fig 5, were represented segmentations corresponding to the minimum, median and maximum distances for all *D*_1_ and *D*_2_ with the constrained spectral clustering *K* = 6. For the C.S. (first column), we obtained *D*_1_ = 1.8 mm, 13.6 mm and 27.0 mm respectively. For the P.O.S it was *D*_2_ = 2.7 mm, 10.7 mm and 31.2 mm.

*Distances between Automatic and Freesurfer segmentation*. Secondly we evaluated another metric to quantify the adequacy between spectral clusters and anatomical information. The rand distance was computed between each automatic segmentation and the corresponding one obtained by Freesurfer. The values offered another look on the variability of the spectral clusters. Figures 6 showed the segmentations with minimal, median and maximal distances.

**Figure 4:**
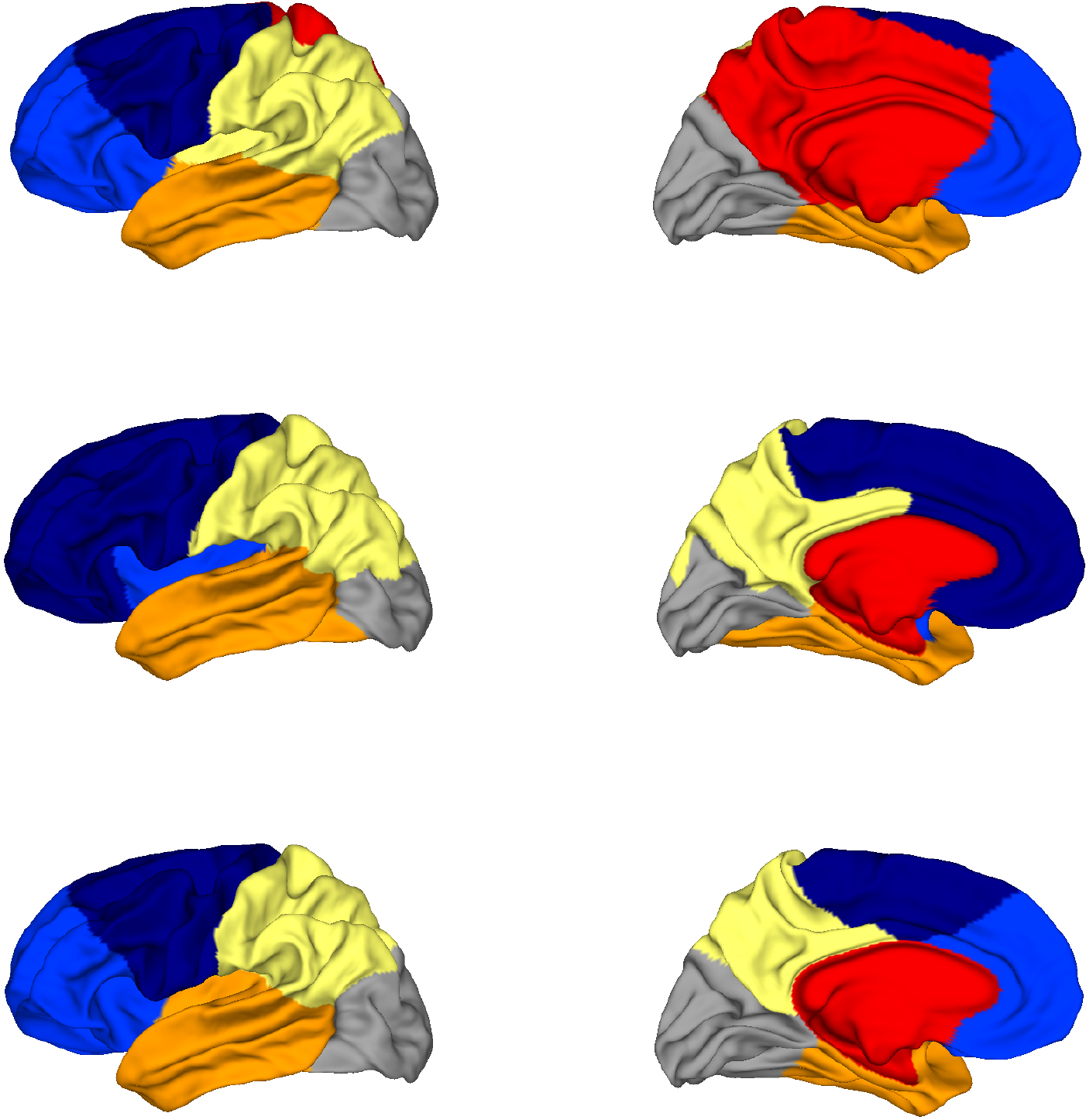
Top row: Consensus segmentation with the unconstrained spectral approach. Middle row: Consensus segmentation of Freesurfer lobes. Bottom row: Consensus segmentation with the constrained spectral approach.

**Figure 5:**
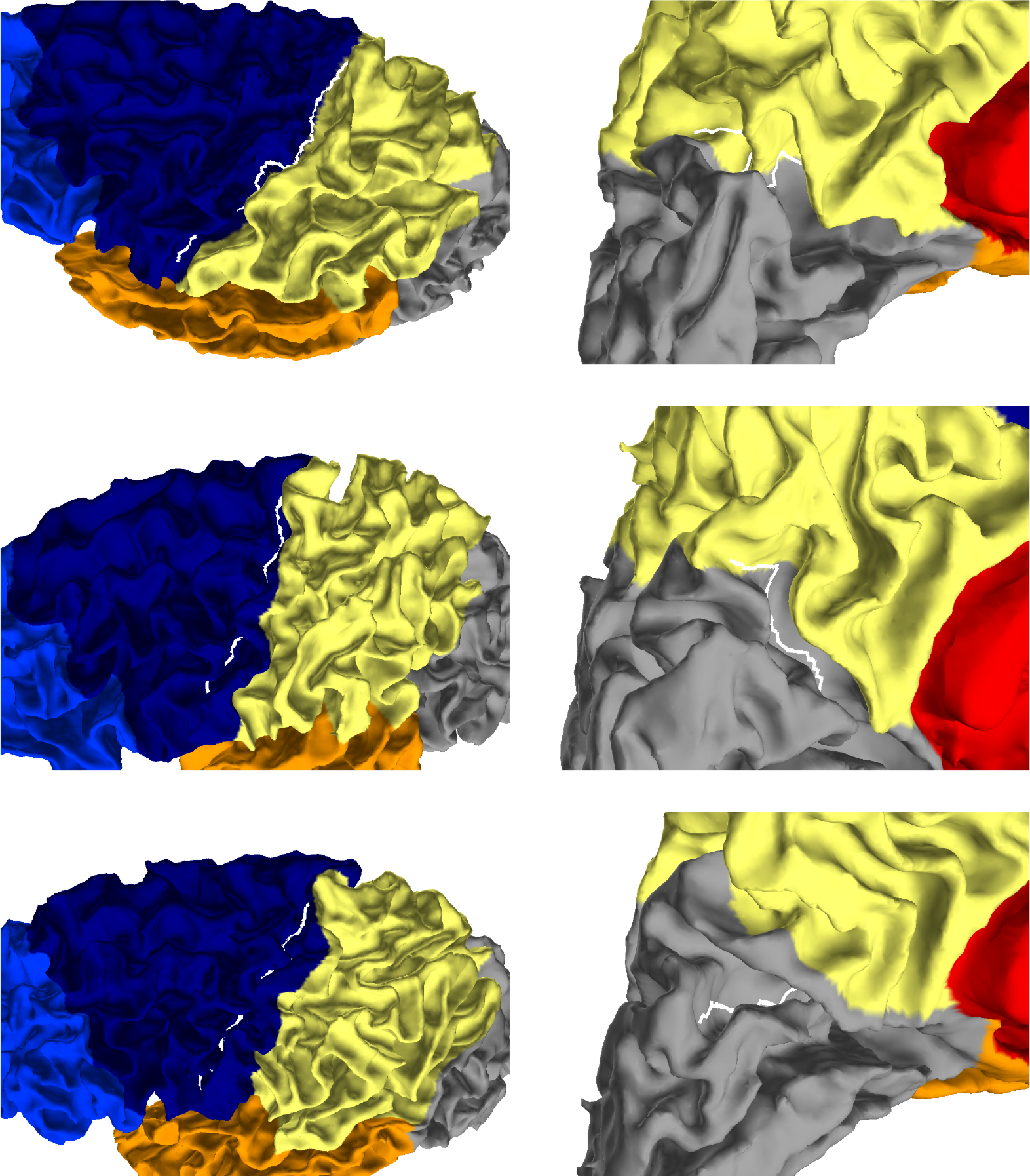
Segmentation boundaries close to central sulcus (left column) and parietooccipital sulcus (right column) with the constrained clustering and *K* = 6. In both cases, the sulcal lines are superimposed in white. Each row corresponds to different positions in the distribution of distances *D*_1_ and *D*_2_ (minimum, median and maximum).

**Figure 6:**
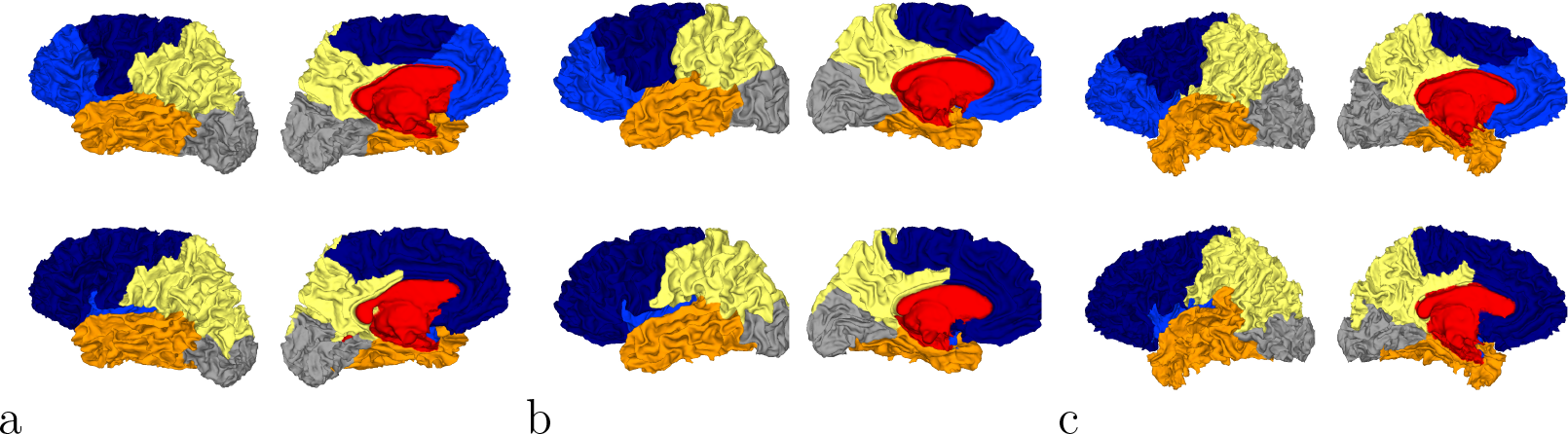
Three spectral segmentatons with constrained cingulate pole (first row) for which the distance with Freesurfer (second row) is the smallest (a), median (b) and the largest
(c).

Quantitatively the values ranged between 0.126 and 0.153 in the constrained case (Fig 6).

*Significance of spectral segmentations*. To go further we obtained, for each subject, 500 random rotations of Freesurfer lobes on the sphere to generate random partitions that share similar area with the initial Freesurfer segmentation. Those random maps were then used to compute random distances with the different spectral segmentations at the individual level. Eventually we were able to compare the partition distance to the randomly generated distributions of distances. We counted how many subjects among the 62 had a p-value below a significance level of 0.01 as obtained from the statistical procedure in 2.5. We summarized the result in Table 1 for the different cases (constrained and unconstrained, *K* = 5, 6 or 7). It confirms that constraining the cingulate pole for *K* = 6 improves the ratio of subjects showing a statistical relationship with brain lobes (from 98.4% to 100%).

**Table 1:** Number of subjects among the 62 whose p-value is less than 0.01. Here *K* = 6 (*K* = 5) in the unconstrained (constrained) case.

| | $K$ ev | $K + 1$ ev |
| --- | --- | --- |
| Unconstrained | 61 | 26 |
| Constrained | 37 | 62 |

*Optimality regarding the number of clusters*. We computed partition distances between constrained spectral approaches with *K* labels and *K* eigenvectors when *K* varies between 3 and 10, given *K* = 2 is pointless (two regions with the cingulate pole and its complement part). On Fig 7 left, we observed the distribution of rand distances and the median values (red) ranged between 0.15 and 0.3 with a clear minimum for *K* = 6. On Fig 7 right, we added information obtained through our random rotations procedure. We displayed both the distribution of individual p values (boxplot) and the ratio of subjects for which the rand distance was significant with respect to the random distribution. This ratio was 1 for *K* = 6 and 0.38 for *K* = 7 (24 subjects). For all the other values, the ratio was below 0.3 suggesting poor association between spectral segmentations and Freesurfer lobes. On Figure 2SI we showed all the consensus segmentation obtained when *K* varied between 3 and 10.

### 3.4. Group analysis of spectral clustering

In this part we further analysed the ability of the spectral clustering approaches to delineate automatically brain lobes. We evaluated first the overlaps between each Freesurfer lobe and the corresponding spectral region and then by comparing the two strategies described in 2.3. Those steps necessitated before a consistent matching between the regions, at the subject and group level, by using the hungarian algorithm.

**Figure 7:**
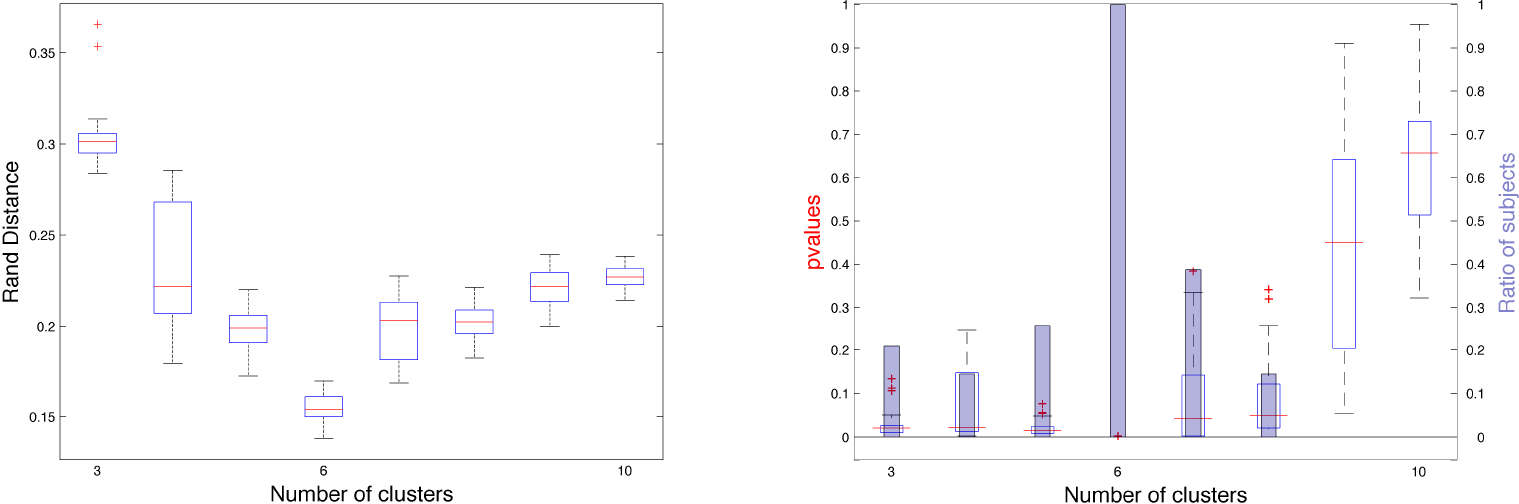
Left: Rand distances between constrained spectral segmentation with *K* regions and Freesurfer segmentation with 6 lobes. Right: p-values for each subject represented as boxplots and proportion of subjects (blue bars) for which the p-value is < 0.01.

*Lobe by lobe statistics*. Beyond the global rand distances shown in S.I., we proposed Dice coefficients between each Freesurfer lobe and the corresponding one in two spectral approaches (unconstrained, constrained, with 6 eigenvectors). The corresponding frontal lobe was obtained by concatenating two spectral regions (light and dark blue in most of the previous figures). The Insula was not included since it had no corresponding region after the grouping strategy for the frontal lobe. The results were summarized in table 2. For all lobes except for the occipital one (almost no change), the mean Dice coefficients were improved by constraining the cingulate pole, particularly for the parietal lobe.

*Comparison of the two group strategies*. In table 3 we showed the 6 Dice coefficients between the regions obtained with the individual spectral clustering and the joint group clustering. We remarked that dice values were large (above 0.93) and close to the perfect overlap (= 1). For all lobes except the temporal one, the overlap was improved when the constraint on the cingular pole was added.

**Table 2:** Dice coefficients between Freesurfer lobes and each of the 5 corresponding regions obtained with the spectral segmentation. Note that the corresponding frontal lobe is obtained by concatenating two spectral regions. The Insula is not included since it has no corresponding region after the grouping strategy in frontal lobe. First row: unconstrained method with 6 eigenvectors. Second row: constrained method with 6 eigenvectors.

| Cingulate | Frontal | Parietal | Temporal | Occipital |
| --- | --- | --- | --- | --- |
| 0.39±0.04 | 0.89±0.01 | 0.66±0.05 | 0.82±0.02 | **0.82±0.03** |
| **0.91±0.02** | **0.94±0.01** | **0.83±0.03** | **0.87±0.01** | 0.81±0.03 |

**Table 3:** Dice coefficients for each of the 6 spectral regions obtained with the individual and the group strategy. First row: unconstrained method with 6 eigenvectors. Second row: constrained method with 6 eigenvectors.

| Front | PreFront | Occip | Pariet | Tempo | Mesial |
| --- | --- | --- | --- | --- | --- |
| 0.95±0.03 | 0.93±0.04 | 0.96±0.03 | 0.94±0.03 | **0.97±0.02** | 0.97±0.02 |
| **0.96±0.02** | **0.98±0.01** | **0.97±0.02** | 0.94±0.03 | 0.96±0.03 | **1.00±0.00** |

On Fig 8 we showed three subjects representing maximal, median and minimal overlap between the two strategies (with constrained cingulate pole) among the group of 62 subjects. For the subject with minimal overlap the two lowest Dice coefficients were 0.88 for both temporal and parietal lobe, values that coincided with the the visual impression (shift of the yellow/orange boundary). Therefore, even if minimal, this overlap corresponded to very good dice coefficients.

**Figure 8:**
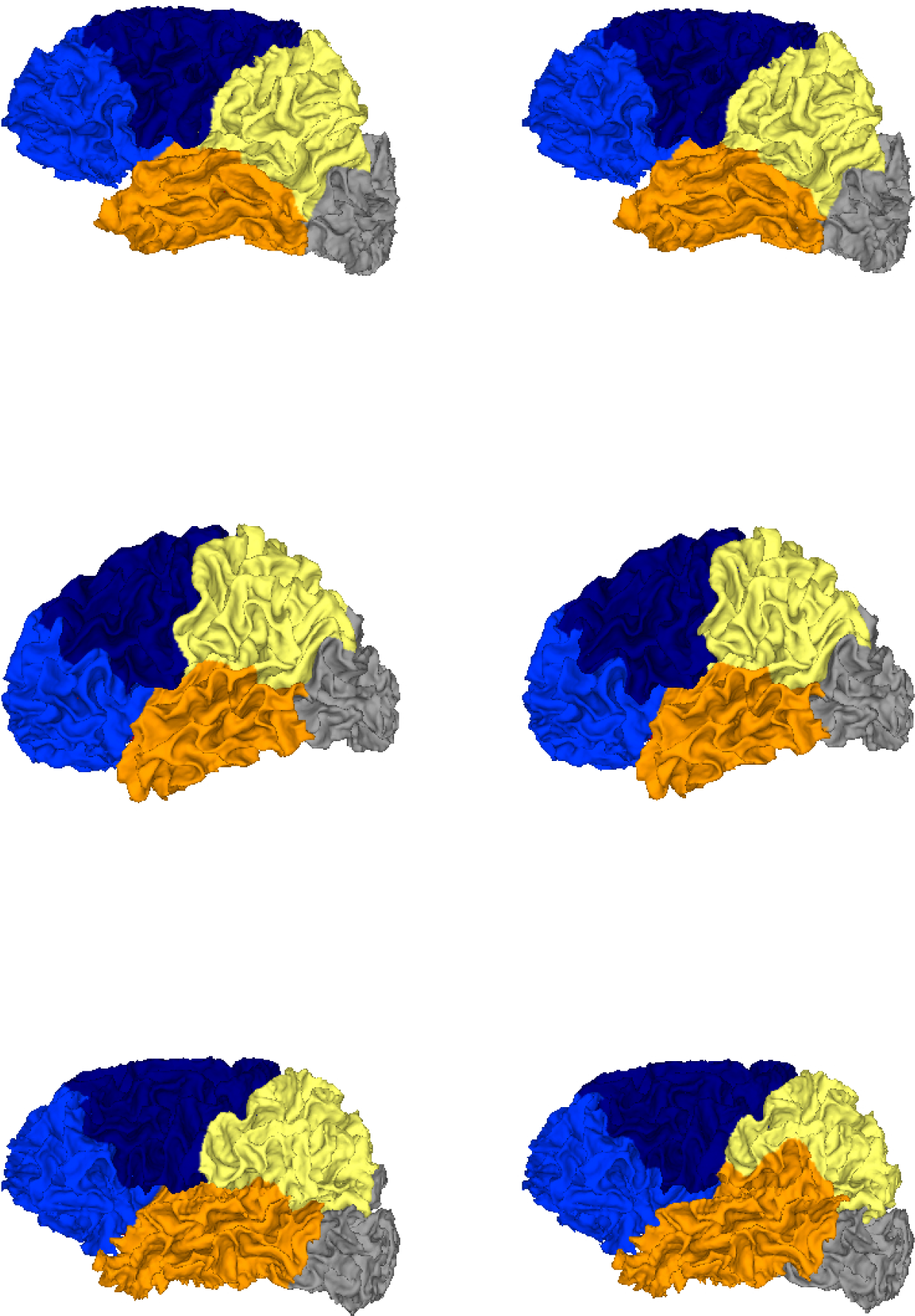
Constrained spectral segmentation for three subjects with individual (left) and group approach (right). From top to bottom, the three subjects correspond to the maximal, median and minimal overlap between the two strategies among the 62 subjects.

### 3.5. Robustness of the constrained spectral segmentation

*Effect of surface smoothing*. For each subject we applied the mean curvature flow to smooth the folded geometry of the cortical meshes. For several iterations of the process (*t* = 1,10, 20, 50,100, 200, 300, 500, 750,1000) we ran the constrained spectral clustering procedure with 6 eigenvectors and computed the partition distance between the resulting segmentation and the initial segmentation. The variability of this distance was illustrated on Fig 9 left. We observed a regular increase of the mean value till an interval [0.04 — 0.09] and a spreading of the distribution as well. Compared to the rand distances found between constrained spectral parcellations and Freesurfer lobes (around 0.15 on Figure 7 or Table 2SI) which reflected good agreements, one can therefore consider that the perturbation of spectral parcels during the smoothing process is really small, meaning that the segmentaion process is rather robust. On Fig 9 right we observed the constrained spectral clustering for a surface whose rand distance correspond to the median value (0.06) of the distance distributions for 1000 iterations. The most visible change can be seen on the mesial face with a slight bifurcation of the yellow/blue boundary that has a counterpart on the external face close to the central sulcus.

*Segmentation of the immature brain*. On Figure 10 we showed the constrained spectral segmentation during early development on 15 fetuses whose gestational age ranges from 21 weeks to 34 weeks. We observed a nice consistency between the different regions across ages. Moreover the position of the parcells was qualitatively similar to what was observed for adult brains. But more precisely the analogous of frontal/parietal boundary crossed the central sulcus which was not the case for adult brains as revealed in particular by Figure 5. It is also possible to quantify the distance between C.S. (respectively P.O.S) and its corresponding boundary, except for the two youngest subjects (21 and 24 weeks gestational age). For the C.S. the absolute values range from 0.7 mm (28 w GA) to 6.3 mm (25 w GA) while for the P.O.S it was from 1.3 mm (25 w GA) to 6.1 mm (33 w GA). In both cases the correlation of those values with age were found non significant (*R* = 0.46 and *R* = -0.07 respectively) when correcting for the increasing global size of the cortical surface, as measured by its longest geodesic (Lefevre et al., 2012).

**Figure 9:**
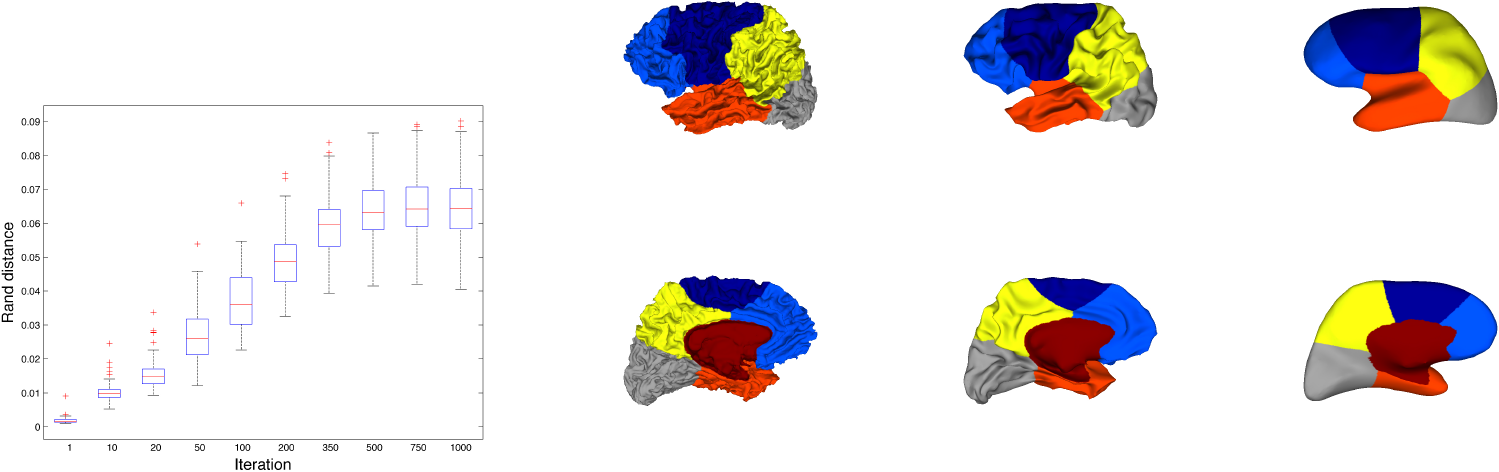
Left: Distribution of rand distances between initial constrained segmentations and segmentations for increasing number of iterations in the smoothing process. Right: illustration of the smoothing process for a surface whose rand distance correspond to the median value (0.06) of the distance distributions for 1000 iterations. 3 iterations are considered (0, 100 and 1000).

## 4. Discussion and Conclusion

The major applicative result of our general Spanol approach is the possibility to segment a cortical surface in a few parcels that have strong similarities with brain lobes. These similarities can be precisely quantified by partition distances and a dedicated statistical procedure has been proposed to determine how far the resulting parcels stood from randomly segmented lobe-sized parcels. The segmentation process can be entirely unsupervised, using only the geometrical properties of the hemispheric mesh shape, as proposed in our preliminary work (Lefevre et al., 2014). However, we improved dramatically the global overlap with the four main lobes (frontal, parietal, temporal and occipital) obtained by Freesurfer software by adding limited constraints only, tagging points that belong to non-cortical regions of the mesh, namely the cingulate pole. This could be seen as a semi-supervised clustering at the level of the whole mesh, but remains strictly unsupervised regarding the cortical level. We would insist on two main neurological relevancies of Spanol, and further discuss interesting methodological considerations raised by its development.

**Figure 10:**
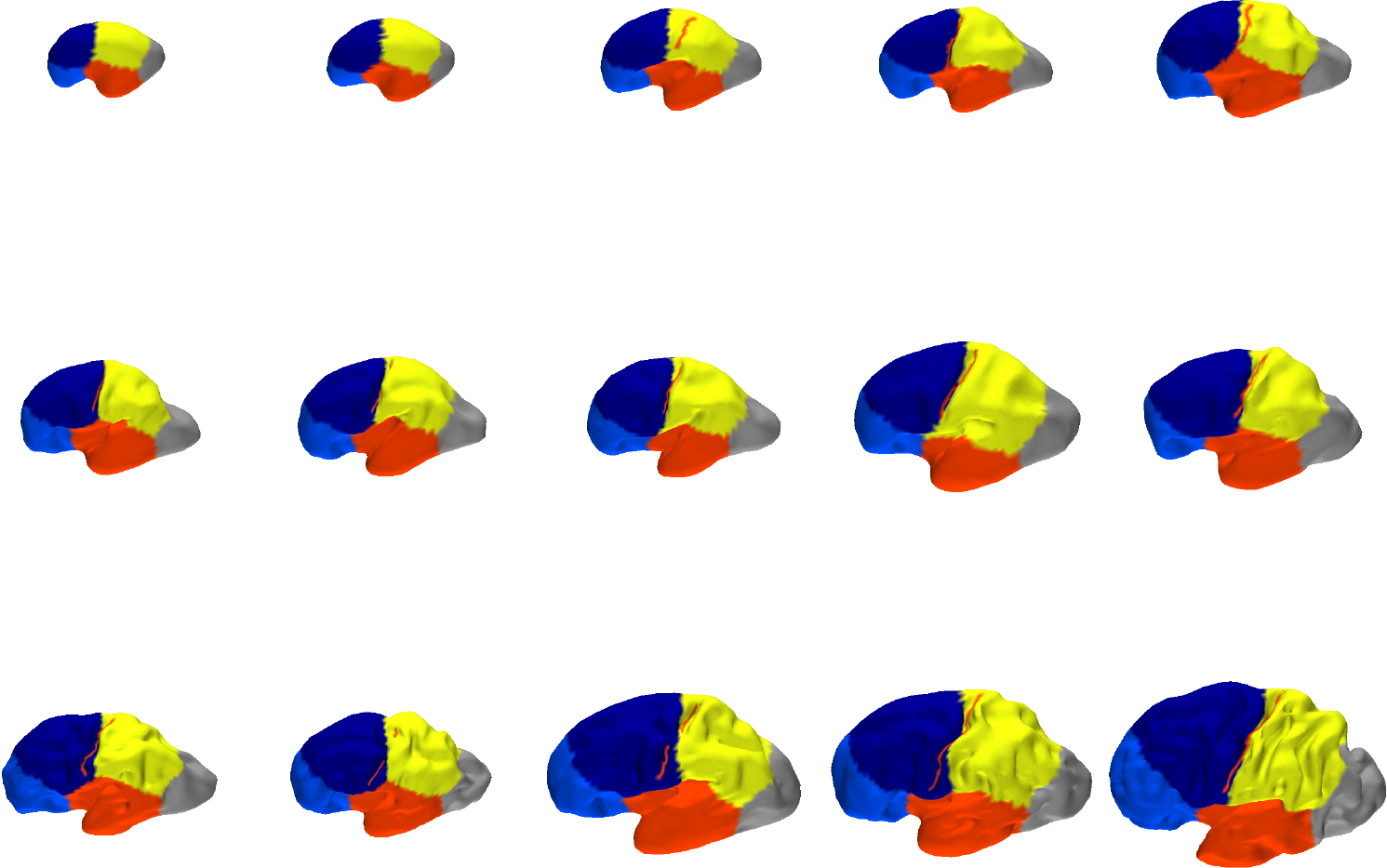
Constrained spectral segmentations of fetal brains with preserved proportions. From left to right, top to bottom: 21, 24, 25, 28, 29 (5), 30, 32, 33 (3), 34 weeks gestational age. The central sulcus, when present, is superimposed in red.

### 4.1. Towards an efficient virtual neuroanatomist

First, to our knowledge it is the first attempt to extract in a fast, simple, reproducible and almost unsupervised manner a parcellation of a cortical surface in lobes. This approach does not require any atlas and can be applied as it is to triangular meshes, at the individual level, in a purely intrinsic way. Although some local improvements can be expected according to experimental needs, they may be achieved with complementary segmentation steps, either supervised or not, following Spanol backbone. For instance, the strict respect of bottom sulci lines along some lobar boundaries (e.g. for C.S. or P.O.S) would be a possible finishing stage, in the same spirit as (Klein and Tourville, 2012). The exclusion of otherwise segmented Insula before spectral clustering has been also tested without disturbing the relevance of lobar boundaries. Indeed, the implementation of Spanol could allow regional analysis of morphometric parameters, in the spirit of (Toro et al., 2009), to be rapidly performed without requiring complicated pipelines and potentially skewing normalisation steps.

### 4.2. A new insight in neurodevelopment

Second and more fundamental, spectral lobes are obtained from the intrinsic geometrical informations contained in the low-frequency eigenfunctions only. Yet, the lobar limits given by Spanol are not only consistent with the classical sulcal ones (C.S., P.O.S) but also for the ill-defined one such as the lateral parietal occipital or the temporal occipital. Even the segmentation of a prefrontal sub-lobar region seems rather consistent. While classical anatomy relies on almost ad hoc landmarks such as the preoccipital notch, or locally defined virtual lines (e.g. occipital temporal limit), spectral analysis kind of intergrade the whole shape into objectively defined boundaries. Indeed, the fact that lobes can be intrinsically defined from the shape of the brain is per se an intriguing result. Interestingly, not all the geometrical information is required, but only the one of the low-frequency eigenfunctions that give mostly information on the global shape of the brain (Seo and Chung, 2011; Germanaud et al., 2012). It may suggest a strong relationship between global shape of the cortical surface and the appearance of great primary sulci that defined main lobar limits (C.S., P.O.S), but also with functional aspects associated with lobes. Anyway, this dependency upon global shape only would explain the stability of spectral segmentation of lobes along development regardless of the full expansion of the main sulci. This claim has been empirically verified by smoothing the cortical surfaces to decrease the influence of sulci and gyri in the geometry and by quantifying the stability of spectral lobes during this deformation.

Moreover the good reproducibility of spectral lobes on a small population of fetuses with various folding complexity is another argument in favor of a very early determination of brain lobes, even from a very smooth cortical surface. This question is also closely intertwined with the emergence of primary folds during fetal development. Among various explanations of this phenomenon, the mecanical hypothesis (Tallinen et al., 2016) is seducing in our case because of oscillatory properties of the surface as proxies of its ability to fold and determine the position of primary folds. Indeed, the question of the relationship between shape and function is all the more interesting since one inverts the classical paradigma and question the fact that the determinent of shape may impact the functional outcome (Foubet and Toro, 2015). So far, it is a very general question, not restricted to the lobar one.

From a more practical point of view, Spanol offers a promising way to define automatically brain lobes in fetuses or newborns because of this nice continuity in the ontogeny. It would remain to test more systematically our method in larger longitudinal databases with complementary information provided by cytoarchitecture, connectivity or functions.

### 4.3. Interesting methodological considerations

Spanol provides a segmentation of cortical surface by using a *K*-means clustering of Laplace-Beltrami operator eigenfunctions. In other terms, only few low frequency descriptors of the brain geometry are required (optimum with 6 in our work) to provide a given number of relevant connected regions on the cortical surface. This method has been used at the individual level and at the group level by concatenating all the eigenfunctions of each subject of the group. A major advantage of the second approach is the ability to obtain directly a consistent labelling of the resulting parcels. In the individual case, it requires to solve the assignment problem to match parcels of different subjects. From a computational point of view the group spectral clustering with 62 meshes of 100 — 150*k* vertices takes approximately 1min on a standard laptop which is comparable to 62 individual *K* means (*K* = 6 in our case). The results showed strong similarities between the segmentations obtained at the individual or group level as measured by the rand distance. This result legitimates the group spectral clustering on our data as a way to segment individual meshes in a consistent, reliable and fast manner. It also suggests a very reproducible pattern in the low frequency eigenfunctions of cortical surfaces which has already been pointed out in several works (Germanaud et al., 2012; Lombaert et al., 2013; Lefèvre and Auzias, 2015). It could be interesting to explore whether a group spectral clustering procedure like our’s could be applied successfully to a larger number of regions. Indeed, in the field of computer graphics the co-segmentations of shapes with spectral approaches have been shown efficient but often tested for limited number of components (Sidi et al., 2011). Conversely studies with other kinds of neuroimaging data (fMRI) suggested that spectral clustering approaches would not be satisfactory for large values of *K* (Thirion et al., 2014).

A novelty in our approach concerns the statistical procedure to test whether a given brain segmentation could be considered as randomly sampled from a null distribution of brain parcellations sharing similar properties. We escaped the intractable (and biologically irrealistic) combinatoric of testing all possible partitions of a triangular mesh by proposing random rotations of a segmentation map onto a reference sphere and computing distances with a reference segmentation. This approach is able to find statistical associations between spatial partitions independently of the partition distance used and of the disparity between the number of clusters in the reference and the automatic segmentations. This procedure takes approximately a few minutes for each subject for 500 random samples. With this statistical tests, we are able to determine which subjects have a spectral parcellation that is significantly close to a reference one. At the end we can find the number of clusters which maximizes the number of significant subjects.

Regarding the supervised information injected in fixing the cingulate pole, we have adopted a strategy where eigenfunctions of the original surface are used in the spectral clustering while excluding the points of the constraint. Something more mathematically natural would have been to consider the eigenfunctions of the Laplace-Beltrami Operator with boundary conditions on the borders of the cingular pole. But our choice was motivated by purely pragmatic considerations regarding unsatisfactory segmentations in the second case. Why the sound mathematical framework of Neuman or Dirichlet boundary conditions has failed and why our ad-hoc treatment of boundaries in the cingulate pole is more successful, remains an interesting question that might even be more general than in our specific context of neuroanatomy. Furthermore the idea of introducing constraints in the spectral clustering could be extended, following previous lines of research in semi-supervised spectral clustering with pairwise association between the points of a dataset (”must-link” and “cannot link”) (Lu and Carreira-Perpinán, 2008) or of a mesh (Sharma et al., 2010). There exist also promising ideas in modifying the Laplace-Beltrami operator with extrinsic geometry information to obtain eigenfunctions that follow more precisely high-frequency features (Andreux et al., 2014). Such strategies could then provide the strict respect of lobar boundaries evoked in subsection 4.1.

As closing remarks, our work illustrates, from the applicative point of view of brain anatomy, one of the fascinating properties of the Laplace-Beltrami Operator. For a long time low-frequency patterns of its first eigenfunctions have been well described by theorems like Courant’s nodal one, which estimates the number of spatial oscillations. Despite a few theoretical results, spectral clustering remains a rather empirical way to combine the geometrical information contained in the eigenfunctions. Still, our results show its ability to unveil structure in datasets such as those in neuroimaging. Besides we rediscovered after (Jin et al., 2005) that varying slightly the number of eigenvectors can improve clustering results. This discrepancy with the idealized view of spectral clustering offers also stimulating theoretical perspectives in the future.

## Acknowledgments

This work has been funded by the French Agency for Research (ANR-12-JS03-001-01, “Modegy”).

